# Visualization of extracellular polymeric substances in *Aspergillus niger* biofilms using fluorescent-labeled lectins

**DOI:** 10.1101/2025.05.12.651018

**Authors:** Aswathy Shailaja, Terri F. Bruce, Patrick Gerard, Julia L. Kerrigan

## Abstract

*Aspergillus niger* is a filamentous fungus that adheres to different substrate surfaces and form biofilms consisting of dense hyphal networks embedded in a self-produced gelatinous matrix composed of extracellular polymeric substance (EPS). The EPS mainly contains exopolysaccharides. The objective of this study was to visualize and assess the different exopolysaccharides structure in this extracellular polymeric substance using a combination of two different fluorescent-labeled lectin probes, *Erythrina cristagalli* (ECA) and *Canavalia ensiformis* (Con A). Each lectin is a unique protein that binds to specific a carbohydrate moiety and is classified based on these terminal sugars to which they are binding. Exopolysaccharides are imaged according to their distinct fluorescence color with the help of confocal laser scanning microscopy (CLSM). The biomass, average thickness, and roughness of biofilms were calculated from the Z-stack images using the computer program COMSTAT. A statistically significant difference was observed in the biomass (μm^3^/μm^2^), average thickness (μm), and roughness of the biofilm treated with the two different lectin probes indicating the presence of a higher amount of galactose and β-1,4 N-acetylgalactosamine (β4GalNAc) when compared to the mannose and glucose moieties in the EPS of *A.niger* biofilm. The combination of two lectin-fluorescent probe method staining should help better characterization of *A. niger* biofilms in terms of their heterogeneity with regards to the EPS production.

## 1. Introduction

In microbial biofilms, the cells are encased by a self-produced protective extracellular polymeric substance (EPS), which consists of different macromolecules including polysaccharides, proteins, nucleic acids, and lipids (Flemming and Wingender 2010; Zarnowski et al. 2014; Flemming et al. 2016; González-Ramírez et al. 2016; Erskine et al. 2018; Kaitlin F Mitchell et al. 2016). The composition of biofilm EPS varies by species within the genus (Breitenbach et al. 2016; Kaitlin F Mitchell et al. 2016; Dominguez et al. 2018). While some EPS components are easily degradable, other components are slowly degradable, and their complete degradation requires a broad range of enzymes (Flemming and Wingender 2010). The EPS is also known as ‘dark matter of biofilms’ because of the occurrence of a large amount of biopolymer and the complexity to analyze them completely (Flemming and Wingender 2010; Flemming et al. 2016). The important features of EPS include the capability to adhere to the substrate, cellular cohesion, ability to provide three-dimensional architecture, ability to provide protection from antimicrobial agents and host immune system (Bridier et al. 2010), and serving as a nutritional resource (Baillie and Douglas 2000; Jin et al. 2005; Flemming and Wingender 2010; Ramage et al. 2012; Rajendran et al. 2013; Zarnowski et al. 2014; Kaitlin F Mitchell et al. 2016; Dominguez et al. 2018; Erskine et al. 2018).

Fungi adapt themselves to grow on surfaces and in porous environments. They form biofilms which are medically, industrially, and geochemically relevant on such surfaces (Bhat 2000; Gutiérrez-Correa and Villena 2003; Villena and Gutiérrez-Correa 2006, 2007; Gorbushina 2007; Gamarra et al. 2010; Ramage et al. 2011, 2012; Tourney and Ngwenya 2014; Flemming et al. 2016; Bao et al. 2018). The ability of fungi to grow on porous environments, enhance ion release, and produce protective EPS, support the development of biofilms. Also, EPS formation enables to development destructive diseases in plants (Harding et al. 2010; Flemming et al. 2016). A variety of fungi are known to contribute to biofilms that exhibit a variety of different compositions and amount of EPS (Hawser and Douglas 1995; Chandra et al. 2001; Kuhn et al. 2002; Jin et al. 2004; Seneviratne et al. 2009; Loussert et al. 2010; Gamarra et al. 2010; Ramage et al. 2011). The extracellular fungal components play a crucial role in helping fungi to adhere and colonize the substrate and other living partners. A high amount of EPS production begins after the cell adhesion to the surface (Breitenbach et al. 2016) Moreover, the establishment and maintenance of biofilm communities depend on the amount of EPS production. The component, concentration, and architecture of EPS help determine the biofilm life and their survival rate. Based on these factors, biofilm morphology can be smooth, rough, filamentous, dense, or have other characteristics.

Polysaccharides are a major biopolymer of the fungal cell wall and EPS architecture (Baillie and Douglas 2000; Flemming and Wingender 2010; Zarnowski et al. 2014; Mitchell et al. 2016; Dominguez et al. 2018). Different kinds of carbohydrates have been found in fungal EPS such as glucose, mannose, galactose, rhamnose, xylose, and fucose (Mahapatra and Banerjee 2013; Breitenbach et al. 2016; Chatterjee and Das 2020). Some of the fungal cell wall polysaccharides are water-soluble and can reach the extracellular spaces (Flemming et al. 2016). Hence, they can migrate from cell proximity and contribute to EPS formation. The soluble α and β-glucans have been identified as major polysaccharides of the biofilm extracellular matrix of several fungi (Ruiz-Herrera 1991; Breitenbach et al. 2016; Kaitlin F. Mitchell et al. 2016; Rodrigues et al. 2018). Also, the major fungal biofilm exopolysaccharides are heteropolysaccharides with cross-linking and branching(Flemming and Wingender 2010). In filamentous fungi, the cell wall and EPS are exclusively composed of different heteropolysaccharides (Bardalaye and Nordin 1976; Beauvais et al. 2014; Kaur and Singh 2014; Mahapatra and Banerjee 2013; Breitenbach et al. 2016). Galactosaminogalactan (GAG) is a heteropolysaccharide found in the cell wall and EPS of several filamentous fungi (Flemming and Wingender 2010; Beauvais et al. 2014; Briard et al. 2016). GAG is a linear and hydrophobic polymer composed of galactose and N-acetylgalactosamine (GalNAc) (Bardalaye and Nordin 1976). This galactose and N-acetylgalactosamine are also present in *Aspergillus niger* biofilms (Bardalaye and Nordin 1976; Mahapatra and Banerjee 2013). In addition, the fungal cell wall polysaccharide mannose and glucose were also reported on *A. niger* biofilms (Villena and Gutiérrez-Correa 2007).

Lectins are a broad class of glycan-binding proteins that have their origin as plant proteins known to agglutinate red blood cells. Generally, lectins are named for the organism from which they are purified, also known as agglutinin (Vectorlabs-Lectins & Glycobiology 2020). Each lectin is a unique protein that has developed to bind specific carbohydrate moieties and are classified based on these terminal sugars to which they are binding. Each lectin will differ in the strength of the binding based on the affinity and avidity for the carbohydrate moiety. Concanavalin A (Con A) developed from *Canavalia ensiformis* (Jack bean) seed selectively binds to α-mannopyranosyl (mannose) and α-glucopyranosyl (glucose) residues (Concanavalin A conjugates -Product information n.d.). *Erythrina cristagalli* (ECA) is purified from *Erythrina cristagalli* (Coral Tree) seeds which have specificity towards galactose and β-1,4 N-acetylgalactosamine residues (Vectorlabs-lectins &Glycobiology 2020).

The objective of this study was to address the heterogeneity and complex composition of EPS by imaging and, measuring different exopolysaccharides in EPS using a combination of two different fluorescent-labeled lectin probes. *Canavalia ensiformis (*Con A)-TRITC is a red fluorescent lectin conjugate with specificity towards mannose and glucose. In contrast, *Erythrina cristagalli* (ECA)-FITC, which is a green fluorescent lectin conjugate that shows specificity towards galactose and β-1,4 N-acetylgalactosamine. Visualization and quantifications were done on *Aspergillus niger* biofilms created to model those that develop under drip flow. Biofilms were grown in a drip flow reactor as it creates a low shear environment that allows for liquid to flow along with the glass coupons, imitating the conditions in industrial and household environments (Goeres et al. 2009). Exopolysaccharides were visualized according to their interaction with specific target lectin conjugates with the help of confocal laser scanning microscopy (CLSM). The CLSM-based lectin fluorescent probe gives a more in-depth knowledge of EPS architecture.

## 2. Materials and Methods

### Fungus strain, culture conditions, and inoculum

*Aspergillus niger* van Tieghem (ATCC 6275) was maintained in petri plates (90mm diam.) containing Sabouraud dextrose agar (SDA, Difco) sealed with parafilm at 21-25°C. From a 7-day-old culture, a spore suspension was prepared, and the spores were dislodged by pipetting sterile distilled water in 1 mL increments in five different places onto the surface of the culture. The petri plate was gently agitated to dislodge the spores, and the spores were transferred to a 50 mL centrifuge tube with phosphate-buffered saline (PBS) maintained at a pH of 7.4 (Electron Microscopy Sciences). The spores were quantified and adjusted with a Neubauer chamber to the concentration, 10^5^ spores/mL. This suspension was used as the inoculum.

### *Aspergillus niger* biofilm formation

Following the reproducible protocol developed by Kerrigan et.al (2021), the biofilms were produced under low shear. To initiate biofilm formation, 10 mL batch medium (sterile 3% wt/vol Sabouraud dextrose broth (SDB)) and 1 mL inoculum was added aseptically to each channel of the drip flow reactor (DFR) containing a glass coupon. The reactor was then sealed and set to a 10° angle backward. This prohibited the fungal structure from clogging the effluent ports. The reactor was left at room temperature (∼23°C) for 48 hrs switching between dark and light every 12 hrs. The reactor was then positioned at a 10° angle forward to prepare the low shear biofilm run and the system was connected to a carboy containing freshly prepared sterile SDB (0.03% wt/vol). The media was then pumped through the reactor using a peristaltic pump (Shenchen-Lab V series) at a rate of 3 mL/min and the effluent port was connected to a vessel for collecting the spent media. The run was continued for 24 hrs under the same light and temperature regime described above.

### EPS visualization using confocal laser scanning microscopy (CSLM)

Following the biofilm formation in DFR, the glass coupons were collected and gently rinsed with phosphate-buffered saline (PBS) (pH 7.2; Electron Microscopy Sciences) for 30 sec without disrupting the biofilms. Transferring glass coupons from the reactor to petri dish can detach an intact biofilm; therefore, extra care was given while lifting and transferring the glass coupons. For the EPS staining, a mixture of two lectins, *Erythrina cristagalli* agglutinin (ECA) tagged with fluorescein isothiocyanate (FITC) (Vector Laboratories, USA) and *Canavalia ensiformis* A (Con A) conjugate labeled with tetramethylrhodamine-isothiocyanate (TRITC) (Molecular Probes-Invitrogen, USA) was used. Con A-FITC solution was prepared at 1 mg/mL in 0.1 M sodium bicarbonate (pH 8.3, Mallinckrodt chemicals; USA). One µL of 5 mg/mL solution of ECA-FITC and 1 µL of 1 mg/mL solution of Con A-TRITC were added to 1 mL DI water for each biofilm samples. The biofilm samples were stained with this freshly prepared working solution and incubated for 90 mins in the dark at 30^0^ C. After the incubation, the stained samples were imaged using Leica TCS SPE CLSM with excitation/emission at 488/515 nm for ECA-FITC stain and excitation/emission at 532/580 nm for Con A-TRITC. The images were captured based on their fluorescent color difference at ACS APO 10x, 0.3 numerical aperture (NA) dry objective. The experiment was conducted four times at two locations, the center and the corner in each biofilm.

### COMSTAT analysis of fungal EPS

With the captured images, the two-dimensional projection of biofilm Z-stacks was performed using Leica SPE software LASX. The three-dimensional image constructions were performed using LASX, the observer experiences the sample displayed from several angles in an animation. Once the images were collected, it was followed by COMSTAT analysis, a computer program written in MATLAB, equipped with an Image Processing toolbox. COMSTAT analyzes the image stacks of biofilm captured by confocal microscopes. Each image stack was thresholded prior to quantification. In COMSAT, thresholding the image stack was performed by applying a fixed threshold value which was determined manually. Once the images were thresholded, the data volume was analyzed. The analysis depends on the variables that were selected and the number of image stacks that were acquired using CLSM. Quantitative structural parameters of the biofilms, such as biomass, maximum thickness, and roughness coefficient were calculated using the COMSTAT program. The biomass (μm^3^/μm^2^) represented the overall volume of all voxels that contain biomass pixels in all images of stacks divided by substratum area of the image stack. The mean thickness (μm) of biofilm provided a measure of the spatial size of the biofilm. Biofilm roughness provided a measure of variations in biofilm thickness and was an indicator of biofilm heterogeneity, dimensionless parameter (Heydorn et al. 2000; Bridier et al. 2010).

### Statistical analysis

Statistical analysis (F-test) was performed using the SAS statistical program, JMP (Pro). Three F-test models were performed on biomass, average thickness, and roughness of EPS labeled with lectin probes, two main effect models (lectin and location), and one interaction effect (lectin-location). The sample number within their lectin were considered as random effects. For examining a significant difference between the lectin and locations, the least square mean student’s t-test was performed. For the entire tests, P values of < 0.05 were considered as statistically significant throughout the analysis.

## 3. Results

### Visualization of exopolysaccharides with labeled lectins

The exopolysaccharides associated with EPS were identified by fluorescent labeled lectins coupled with confocal microscopy. The visualization was performed both at the center and the corner portion of the biofilm to understand the variation between the two locations (Figs.1.1 and 1.2). As seen in the images, the exopolysaccharide distribution is not homogenous (Figs.1.1 and 1.2). It was also observed that the composition of ECA binding carbohydrates (galactose, and β-1,4 N-acetylgalactosamine) and Con A binding (mannose, and glucose) carbohydrates are varying in the center and corner portions. From these images, it was observed that the composition of galactose and β-1,4 N-acetylgalactosamine (β4GalNAc) is higher than mannose and glucose moiety in the EPS of *A. niger* biofilm.

**Fig. 1.1.**
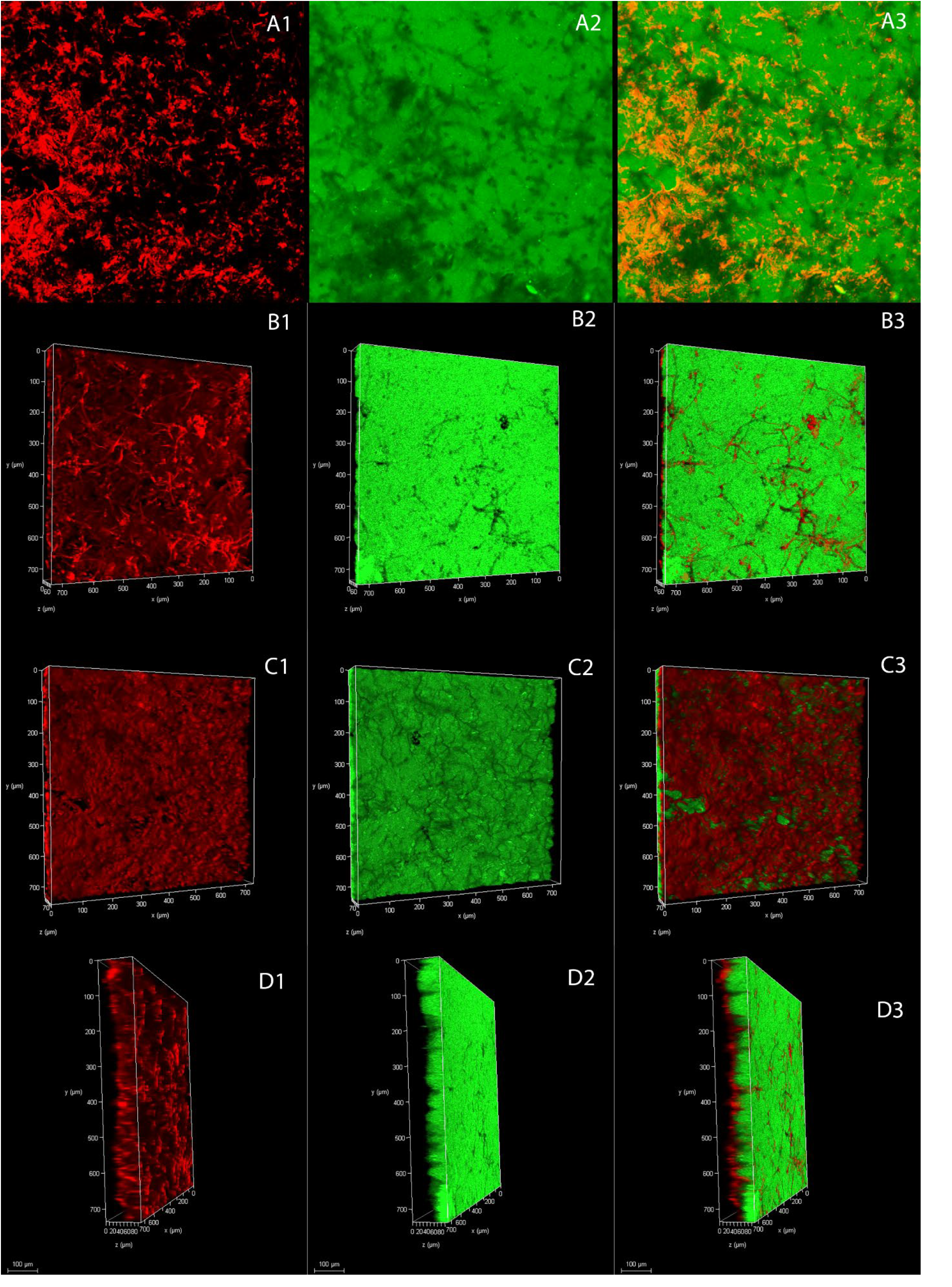
CLSM images of *Aspergillus niger* biofilm grown on a glass coupon (center portion). EPS stained with combination of ECA-FITC and Con A-TRITC. **(A1, B1, C1, and D1)** mannose and glucose moieties fluoresced red with Con A-TRITC, **(A2, B2, C2, and D2)** galactose, and β-1,4 N-acetylgalactosamine (β4GalNAc) moieties fluoresced green with ECA-FITC, **(A3, B3, C3, and D3)** overlay of carbohydrate moieties. **(B1-B3)** Surface 3D view of EPS labeled with ECA-FITC and Con A-TRITC. **(C1-C3)** Bottom (attached to glass substrate) 3D view of EPS labeled with ECA-FITC and Con A-TRITC. **(D1-D3)** The lateral 3D view of EPS labeled with ECA-FITC and Con A-TRITC, the top layer has relatively higher amount of galactose, and β4GalNAc moieties, fluoresced with ECA-FITC and bottom layer has less amount of mannose and glucose, fluoresced red with Con A-TRITC. Scale bars = 100 µm.

**Fig. 1.2.**
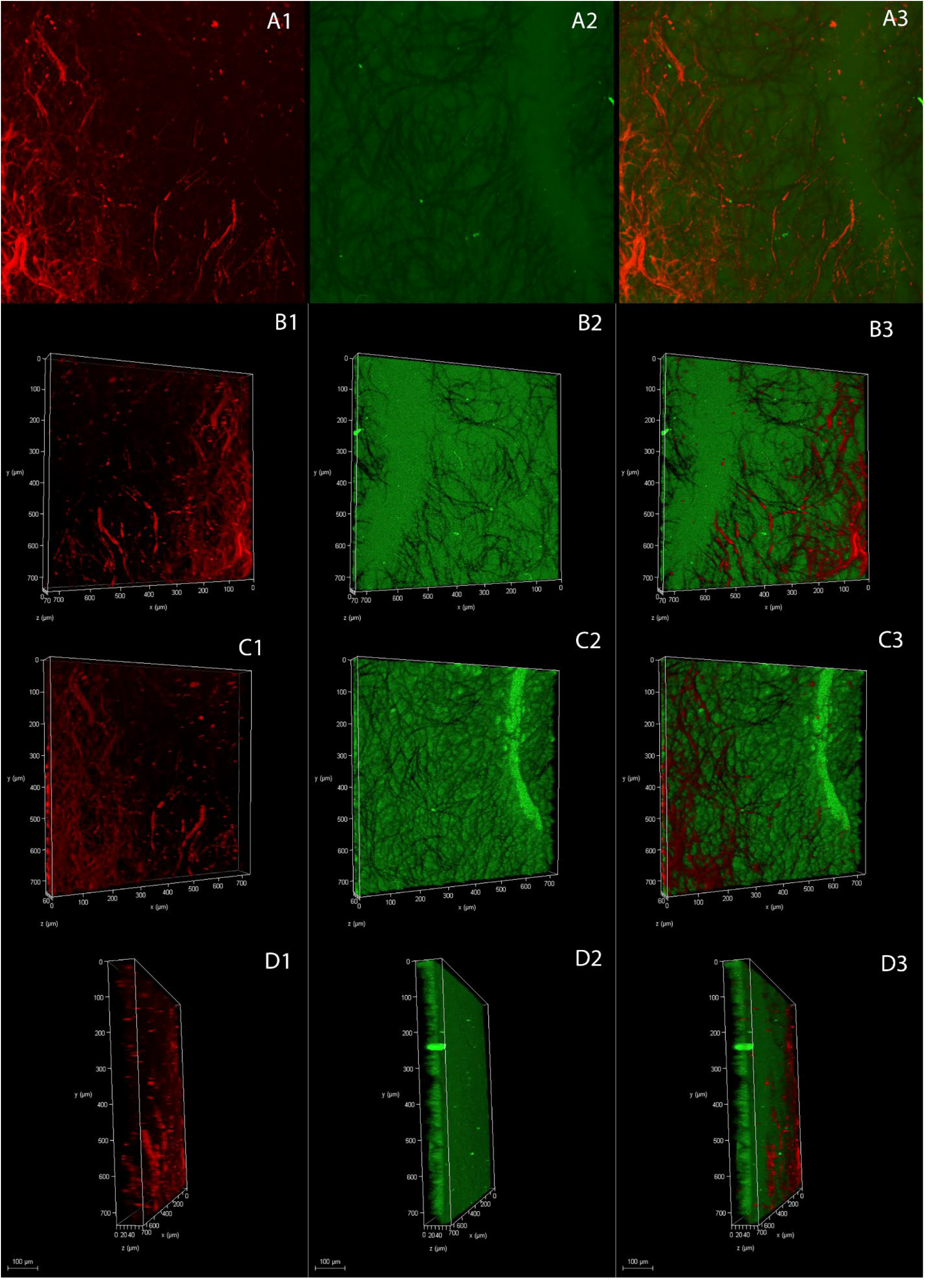
CLSM images of *Aspergillus niger* biofilm grown on a glass coupon (corner portion). EPS stained with combination of ECA-FITC and Con A-TRITC. **(A1, B1, C1, and D1)** mannose and glucose moieties fluoresced red with Con A-TRITC, **(A2, B2, C2, and D2)** galactose, and β-1,4 N-acetylgalactosamine (β4GalNAc) moieties fluoresced green with ECA-FITC, **(A3, B3, C3, and D3)** overlay of carbohydrate moieties. **(B1-B3)** Surface 3D view of EPS labeled with ECA-FITC and Con A-TRITC. **(C1-C3)** Bottom (attached to glass substrate) 3D view of EPS labeled with ECA-FITC and Con A-TRITC. **(D1-D3)** The lateral 3D view of EPS labeled with ECA-FITC and Con A-TRITC, the top layer has relatively higher amount of galactose, and β4GalNAc moieties, fluoresced with ECA-FITC and bottom layer has less amount of mannose and glucose, fluoresced red with Con A-TRITC. Scale bars = 100 µm.

Figures 1.1 (B1-B3) and 1.2 (B1-B3) show the top view of the biofilm on a glass coupon. These figures indicate the presence of higher amounts of galactose and β-1,4 N-acetylgalactosamine when compared to mannose and glucose. Both mannose and glucose are embedded inside the galactose, and β-1,4 N-acetylgalactosamine (Figs.1.1 (A3) and 1.2 (A3). In contrast, in the case of figures 1.1 (C1-C3) and 1.2 (C1-C3), showing the bottom view of the biofilm on a glass coupon, the mannose and glucose were more visible when compared to the top view. It was also observed that, the center portion had a higher concentration of mannose and glucose when compared to the corner portion as seen in figures 1.1 C1-C3.

Figures 1.1 (D3) and 1.2 (D3) show the lateral view of the biofilm on a glass coupon. From these images, the top layer has a higher amount of galactose and β-1,4 N-acetylgalactosamine and the mannose and glucose embedded within them. Interestingly, distinct staining patterns are exhibited by ECA and Con A. The ECA visualized the EPS either in between the *A. niger* cells or cell surfaces. On the other hand, Con A highlighted the hyphal structure with a red ribbon appearance. Con A also stained the EPS surface to a lesser extent than ECA. This difference in the staining pattern is because of the difference in the binding avidity between ECA and Con A. When ECA has an avidity for galactose and β-1,4 N-acetylgalactosamine residues, Con A binds specifically to mannose and glucose residues. This distinct specificity of the two different probes makes it possible to qualitatively examine the heterogeneous cell wall structure/components and, EPS production in biofilms.

COMSTAT analysis was performed to calculate the biomass (μm^3^/μm^2^), average thickness (μm), and roughness from these images. The parameters obtained from this analysis were then compared for further statistical analysis. Three F-test models were used to compare the biomass, average thickness, and roughness of the EPS visualization, two for the main effect (lectin and location), and the other one for the interaction effect (lectin-location). Based on the F-test of biomass, it was observed that there was a significant difference in the two main and interaction effects model (Fig 2.1). The F-test used for lectin in biomass was observed to have a P value of < 0.0001, meeting the criteria of P < 0.05 making the biomass of each lectin statistically different. Moreover, a significant difference between the biomass of the two locations, center and corner of the biofilm was recorded at a value of P = 0.0013. Also, the F-test performed for biomass in the interaction was observed to have a P value of 0.0010, making it statistically significant

**Fig. 2.1.**
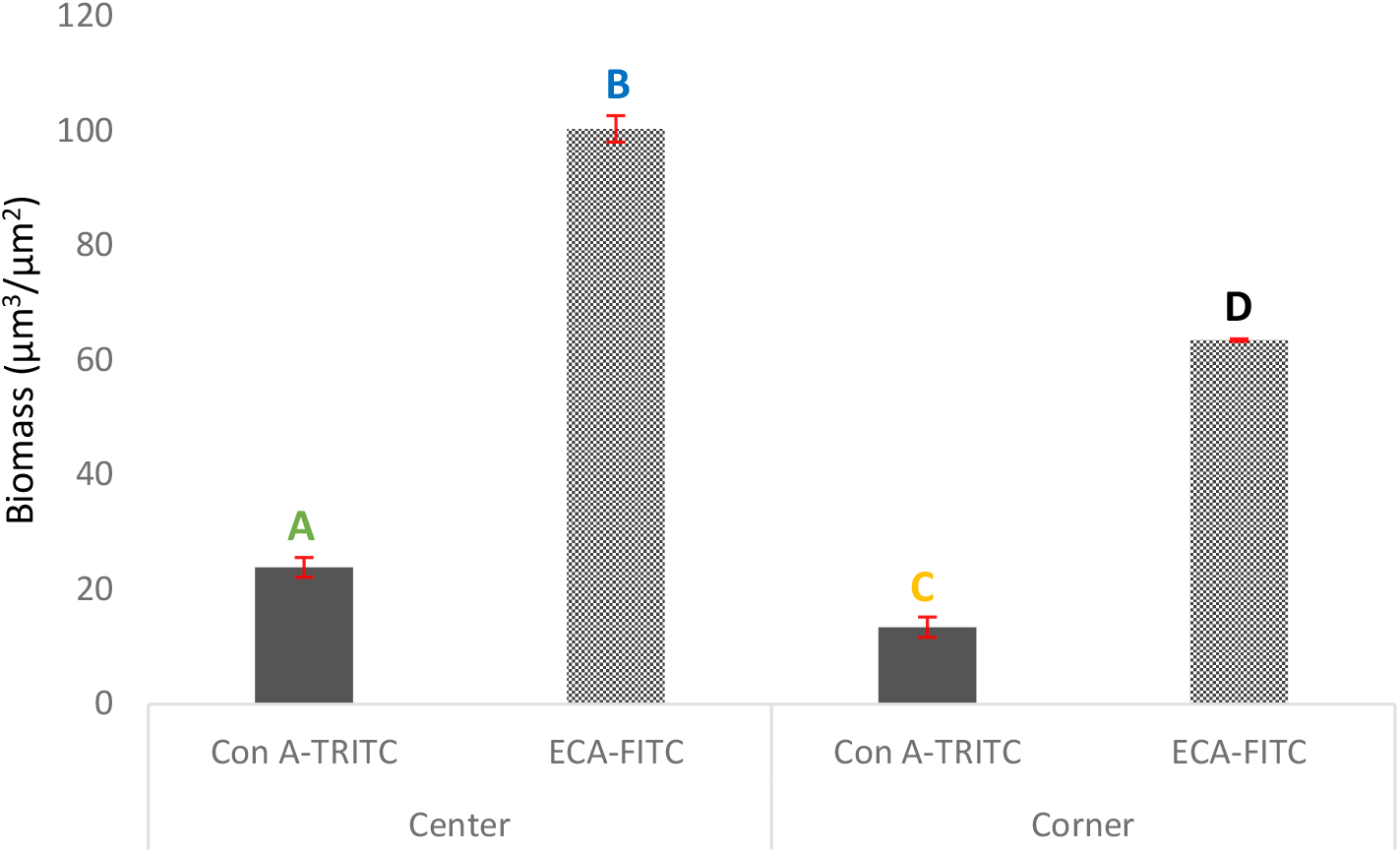
The biomass analysis of *Aspergillus niger* biofilm EPS by lectin-location. Bars represent the least square mean value, and each error bar is constructed using one standard error from the mean. Bars represented by different letters are statistically different. Biomass calculated using COMSTAT. Con A-TRITC binds mannose and glucose residue, whereas ECA-FITC binds galactose and β-1,4 N-acetylgalactosamine.

**Fig. 2.2.**
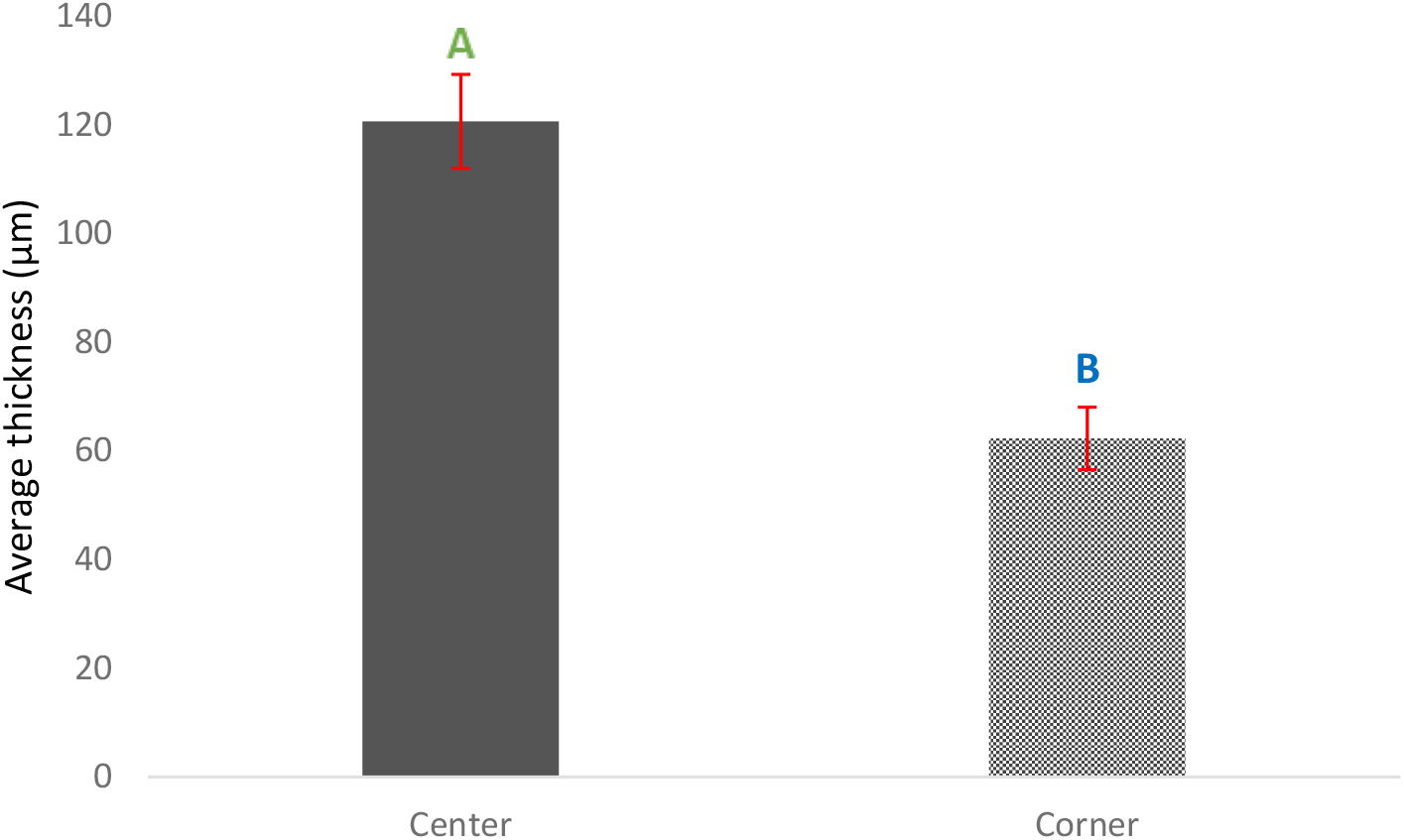
The average thickness analysis of *Aspergillus niger* biofilm EPS by location. Bars represent the least square mean value, and each error bar is constructed using one standard error from the mean. The average thickness of each location is statistically different from one another as represented by different letters. Average thickness calculated using COMSTAT. Con A-TRITC binds mannose and glucose residue, whereas ECA-FITC binds galactose and β-1,4 N-acetylgalactosamine

**Fig. 2.3.**
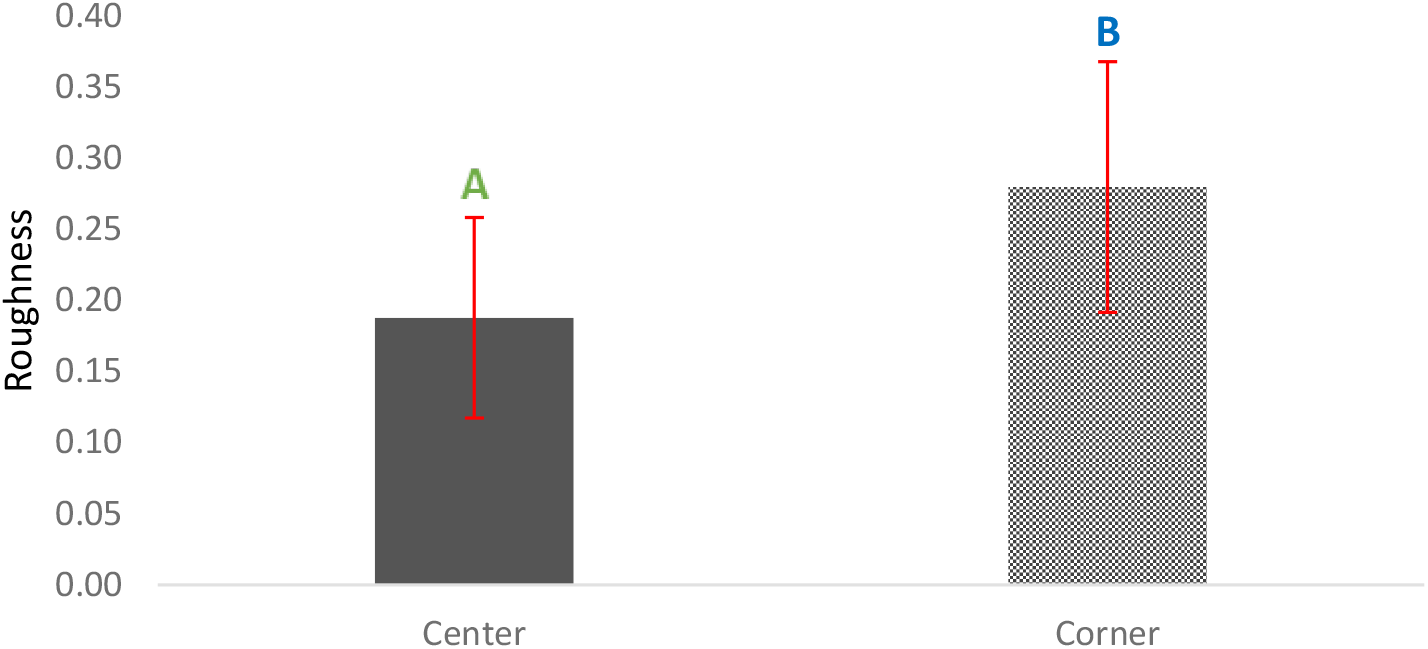
The heterogeneity (roughness) analysis of *Aspergillus niger* biofilm EPS by location. Bars represent the least square mean value, and each error bar is constructed using one standard error from the mean. The heterogeneity (roughness) of each location is statistically different from one another as represented by different letters. Heterogeneity (roughness) calculated using COMSTAT. Con A-TRITC binds mannose and glucose residue, whereas ECA-FITC binds galactose and β-1,4 N-acetylgalactosamine.

Since the P-value from F-test came out as < 0.05, for further effect details of biomass, a student’s t-test was performed. The student’s t-test helps in understanding if there are any significant differences in the biomass among the lectin location interactions (Fig 2.1). Based on the results from the student’s t-test, a significant difference in lectin with respect to its location was observed. This indicates that the impact of the biomass of two lectins compared to each other was not consistent for the center and the corner. Additionally, based on the F-test for average thickness, it was observed that there was a significant difference in the location effect. This implies that the average thickness of the EPS is different between the center and the corner portion. Based on the F-test of average thickness, it was observed that there was a significant difference in the location effect at a value of P = 0.0155, meeting the criteria of P < 0.05. Moreover, the F-test of roughness performed for the location effect was observed to have a value of P = 0.0472, meeting the criteria of P < 0.05 making it statistically significant. This indicates that the homogeneity of EPS is not uniform in the center and corner portion.

## 4. Discussion

In this study, ECA labeled with FITC, and Con A labeled with TRITC were used for the purpose of EPS visualization. ECA is a 54,000 dalton, 2 subunit glycol protein that has specificity towards galactose and β-1,4 N-acetylgalactosamine (β4GalNAc) sugar moiety, whereas Con A-TRIC is a lectin with 104,000 dalton tetramer which binds specifically to mannose and glucose. These two labeled lectins were used because of their specificity for polysaccharides and the two different fluorescent probes which mark the polysaccharides separately. The combination of these two lectins can visualize more polysaccharides in EPS when used independently.

Figures 1.1 (B3, C3) and 1.2 (B3, C3) reveal uneven EPS distribution. In figures 1.1 and 1.2, we can observe that in places where the hyphal structure is present, both the glucose and mannose can be seen. This indicates that the structural component of the fungal cell wall is also a part of EPS production. The combined use of two different lectins with two different fluorescent probes permits the characterization and visualization of *A. niger* EPS. This can be utilized to understand biofilm heterogeneity in terms of EPS.

As seen in figures 1.1 and 1.2, the EPS of the biofilm is not homogenous. Most of the EPS contained galactose and β-1,4 N-acetylgalactosamine, and a smaller contained mannose and glucose. This same trend was observed both at the center and the corner portions of the biofilm. One possible explanation for this phenomenon is that during biofilm formation, a hyphal network was formed. The outer layer of hyphae contained galactosaminogalactan (GAG) that composed galactose and β-1,4 N-acetylgalactosamine. From the previous studies, we observed that the center portion of the biofilm had more active hyphae when compared to the corner portion implying the presence of a higher amount of GAG at the center. Since it was for the first time the EPS was labeled in *A. niger* biofilm using lectin, further studies are required for understanding more about the EPS and exopolysaccharide composition and their ratio. Using multiple lectin conjugates, the carbohydrate composition of the EPS can be quantified. This helps in selecting an anti-microbial agent targeting a specific exopolysaccharide in the EPS.

## 5. Figures and graphs

